# The deep and slow breathing characterizing rest favors brain respiratory-drive

**DOI:** 10.1101/2020.07.30.226563

**Authors:** Baptiste Girin, Maxime Juventin, Samuel Garcia, Laura Lefèvre, Corine Amat, Nicolas Fourcaud-Trocmé, Nathalie Buonviso

## Abstract

A respiration-locked activity in the olfactory brain, mainly originating in the mechano-sensitivity of olfactory sensory neurons to air pressure, propagates from the olfactory bulb to the rest of the brain. Interestingly, changes in nasal airflow rate result in reorganization of olfactory bulb response. Therefore, if the respiratory drive of the brain originates in nasal airflow movements, then it should vary with respiration dynamics that occur spontaneously during natural conditions. We took advantage of the spontaneous variations of respiration dynamics during the different waking and sleep states to explore respiratory drive in various brain regions. We analyzed their local field potential activity relative to respiratory signal. We showed that respiration regime was state-specific, and that quiet waking was the only vigilance state during which all the recorded structures can be respiration-driven whatever the respiration frequency. We used a CO_2_-enriched air to change the respiratory regime associated to each state and, using a respiratory cycle-by-cycle analysis, we evidenced that the large and strong brain entrainment during quiet waking was the consequence of its associated respiration regime consisting in an optimal trade-off between deepness and duration of inspiration. These results show for the first time that changes in respiration regime alter the cortical dynamics and that the respiratory regime associated with rest is optimal for respiration to drive the brain.

## INTRODUCTION

In the movement of the theory of communication through coherence^1^, a new literature sets the respiratory rhythm as a global rhythm, potentially influencing behavior by modulating cortical neurodynamics^2,3^. Indeed, number of studies have described breathing as modulating complex behaviors. In animals, respiration interacts with different functions such as vocalizations, whisking, or licking^4,5^. In humans, a clear relationship exists between breathing and cognitive processing (for review, see Heck et al., 2017). Respiration modulates olfactory memory consolidation^6^ and cognitive performance during the retrieval process^7^. Pain processing is gated during exhalation^8^. In experiments with forced breathing, the gripping force is greater during exhalation than during inhalation^9^. The recognition of a fear expression is faster if the stimulus is presented during the inspiratory phase^10^. Even when no instructions for inspiration are given, subjects inhale spontaneously at the beginning of a cognitive task and these inhalations induce a change in functional connectivity in the brain networks^11^. More interestingly, this last study emphasized the involvement of the nasal respiration by reporting that nasal inspiration induces an increase in performance in a visuo-spatial task.

In parallel, many recent publications show that respiration can modulate ongoing activity of very different cerebral regions both in rodents^12–17^ and humans^10,18^. Most of these studies concluded that respiratory entrainment is dependent on nasal respiration because it is suppressed when nasal airflow is by-passed. In rodents, respiration-locked activity is eliminated in the barrel cortex when nasal airflow is stopped by olfactory bulb removal^12^; in hippocampus, respiration-related slow rhythm is suppressed under tracheotomy^19,20^ and sharp wave ripples probability is no longer respiration-modulated when olfactory bulb activity is inhibited^15^. In human, oscillatory power in olfactory cortex and limbic structures dissipates when the subjects breathe through the mouth^10^. Even if the intervention of a central component cannot be excluded^21^, the main cause of this respiratory entrainment during nasal breathing is likely to be found in the olfactory receptor cells mechanosensitivity to pressure changes in the nasal cavity during successive inspirations and expirations^22^.

A respiration-locked activity propagates from the olfactory epithelium to the olfactory bulb (OB) where it has been extensively described. In brief, respiration has been shown to modulate OB local field potentials (LFPs) fast oscillations^23^, and both unitary discharges^24–26^ and membrane potential of principal neurons^27,28^. Interestingly, changes in airflow rate in the normal range are enough to substantially reorganize the response of the OB: cellular odor-evoked activities and LFP oscillations are strongly modified by nasal flow rate^29–32^. A low/high flow rate favors a beta/gamma regime in the OB, respectively^31^. Combining calcium imaging and an artificial sniffing system, Oka et al. (2009) showed that nasal airflow rate, but not respiratory frequency, is a key factor that regulates olfactory sensitivity of the OB glomeruli.

Yet, nasal airflow varies in both flowrate and frequency according to respiration dynamics. Therefore, if the respiratory drive of the brain originates in nasal airflow movements, then it should vary with respiration dynamics that occur spontaneously during natural conditions. To our knowledge, no study has been conducted to answer this question and the present work aimed at bridging this gap. We took advantage of the spontaneous variations of respiration dynamics during the different waking and sleep states to explore respiratory drive in various brain regions: olfactory bulb, piriform, somatosensory and visual cortices, CA1 and DG hippocampal areas. We analyzed their LFP activity relative to respiratory signal. Unlike how it has been proceeded so far in previous studies^17^, we did not restrict our study to epochs of respiratory steady state. On the contrary, we analyzed signals from complete waking and sleep periods, encompassing the whole range of respiratory frequencies expressed in the freely moving rat. This allowed us to show that respiration regime was state-specific, and that quiet wakening was the only vigilance state where both olfactory and non-olfactory structures can be respiration-driven whatever the respiration frequency (0.8-5Hz) expressed during this state. We used a CO_2_-enriched air in the plethysmograph to change the respiratory regime associated to each state and, using a respiratory cycle-by-cycle analysis, we evidenced that the large and strong brain entrainment during quiet wakening was the consequence of its specific respiration regime consisting in an optimal trade-off between deepness and duration of inspiration. These results show for the first time that change in respiration regime alter the cortical dynamics and that the respiratory regime associated with rest is optimal for respiration to drive the brain.

## RESULTS

We analyzed LFPs and respiration signal, through a whole-body plethysmograph, in 13 rats during active exploration (AE), quiet waking (QW), slow-wave sleep (SWS) and rapid-eye movements (REM) episodes scored as mentioned in Methods. **Figure 1** shows a representative example of simultaneous recordings of respiration and LFPs from anterior piriform cortex (AP), hippocampus CA1, and primary visual cortex (V1) during a 25min-long period where different states alternate (AE red, QW green, SWS blue, REM pink). Typically, respiratory frequency was higher during awake states, with a great variability whereas it was low and with much less variability during sleep states. LFP activities varied both on the low (0.5-12 Hz) and high (20-100 Hz) frequency bands. A major innovation of our study is that we analyzed data on the whole range of respiration frequencies, neither selecting epochs where respiration frequency did not overlap with theta frequency nor focusing on epochs where it was stationary, as proceeded in a similar study^17^. This allowed us to track faithfully respiratory brain entrainment under non-stationary respiration regimes. We first asked to what extent respiration regime varied with behavioral state. For this purpose, we analyzed features of interest, represented in **Figure 2B:**

- instantaneous respiratory frequency (Freq) which was extracted from inhalation and exhalation durations (iDur, eDur)
- inspiration and expiration peak flowrate (iPF, ePF), representing the maximal flow rate, expressed in mL.s^−1^.

**Figure 1:**
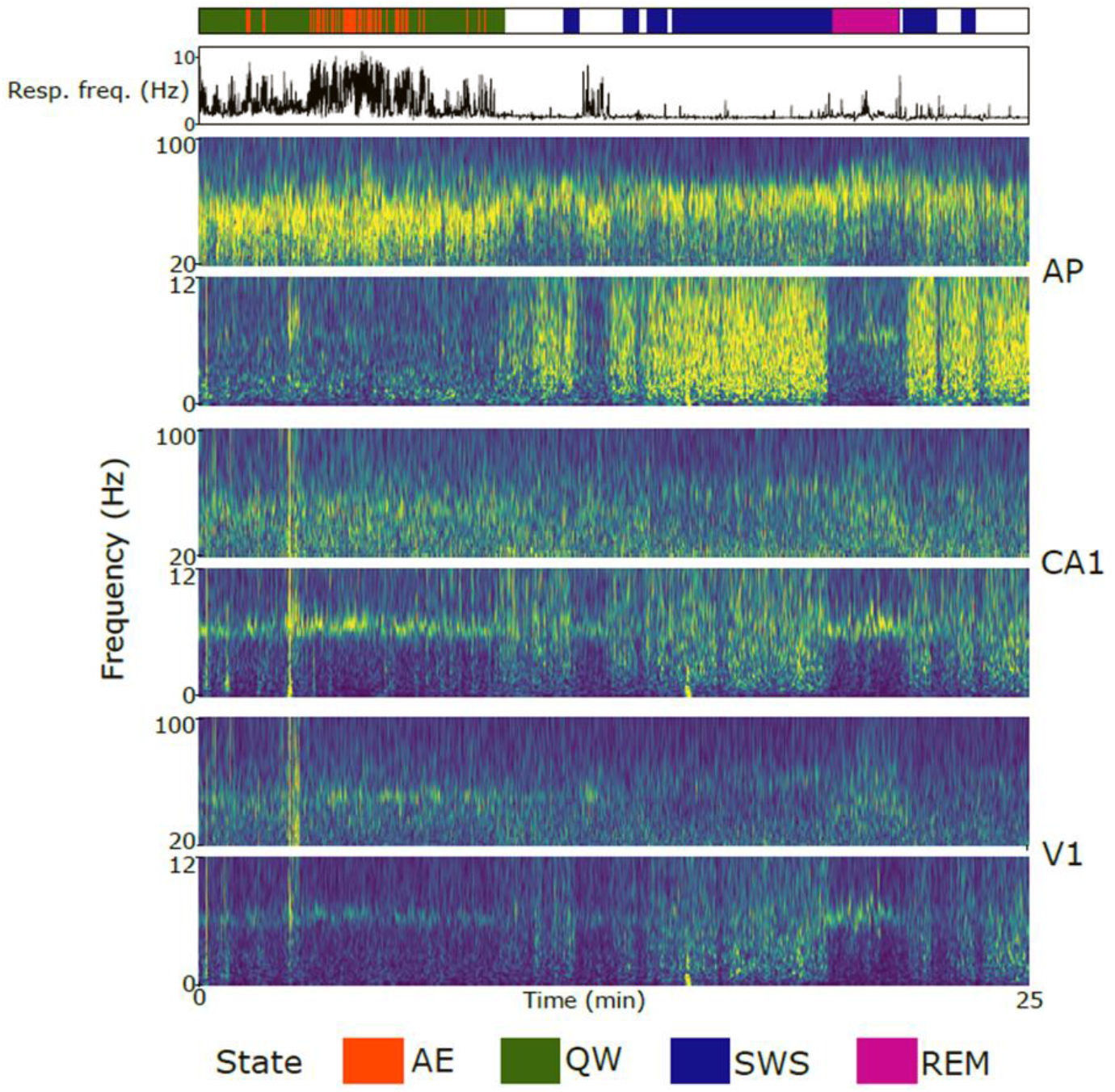
Example of a 25min long recording in anterior piriform cortex (AP), hippocampus CA1 and primary visual cortex (V1) regions. Top panel: states scoring with active exploration (AE, red), quiet awaking (QW, green), slow-wave sleep (SWS, blue), and rapid-eye movements sleep (REM, pink). Periods not scored (not all criteria met, see Methods) appear in white. Second panel: Instantaneous respiratory frequency (in Hz). Other panels from top to bottom: For each structure (AP, CA1, V1), time-frequency representation of LFP in the low (0.5-12Hz) and high (20-100 Hz) frequency range.

**Figure 2:**
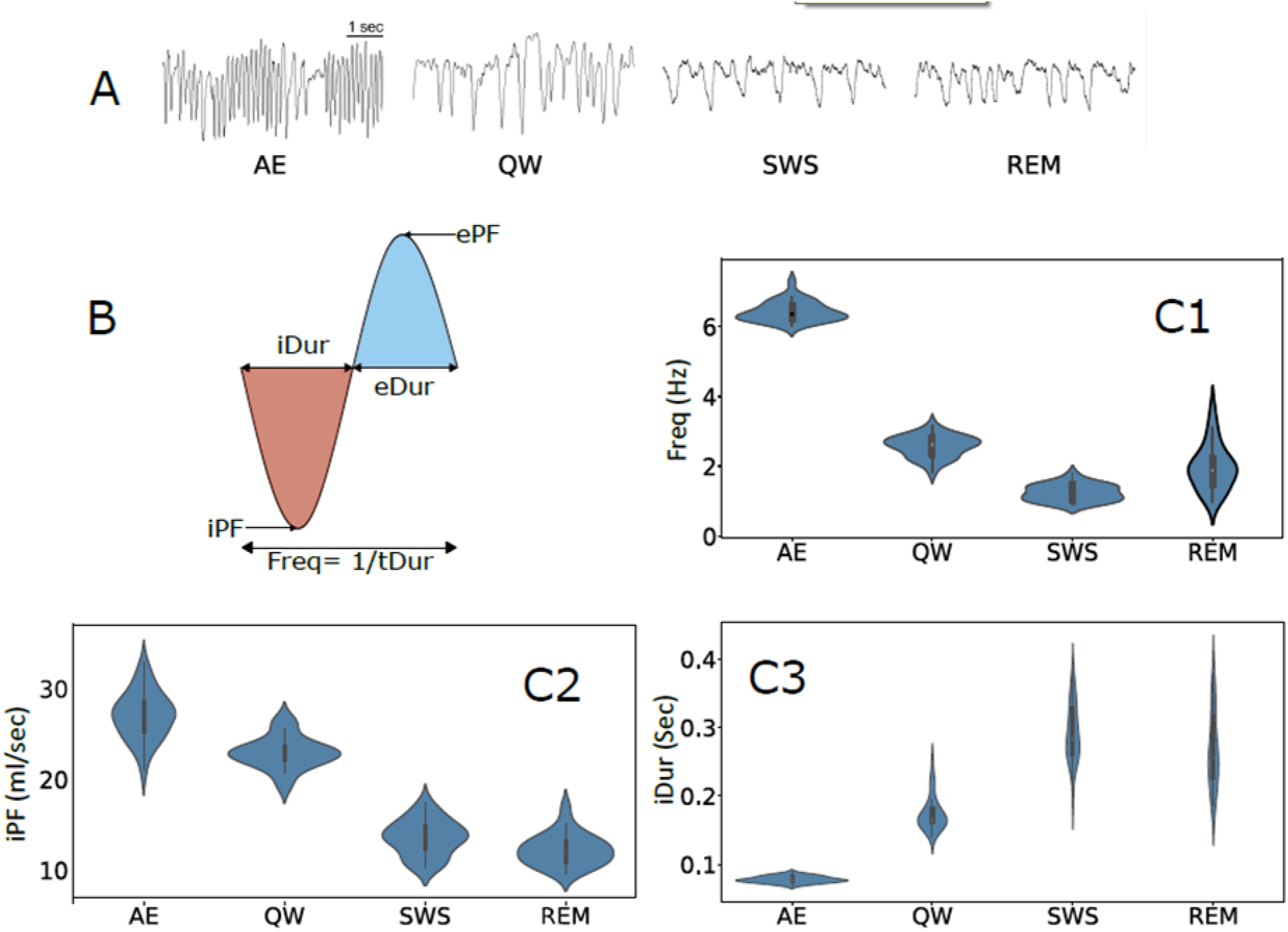
Respiration regimes associated with brain states. **A**: Examples of respiration raw traces during active exploration (AE), quiet waking (QW), slow-wave sleep (SWS) and rapid eye movements sleep (REM). **B**: schematic representation of measured respiratory features. **C**: violin plots of respiration features measured during each brain state: frequency (Freq, C1), inspiration peak flowrate (iPF, C2), and inspiration duration (iDur, C3). One-way ANOVA Test (p<0.05) was first applied for global comparisons and then posthoc Tuckey (HSD) test for multiple comparisons. For all the comparisons between states and for each feature, p<0.001 except iPF for REM *vs* SWS p=0.082 and iDur REM *vs* iPF p=0.063. See sample size (number of rats) in Table 1.

Representative raw traces in Fig.2A illustrate respiration variations between the different states, which are confirmed by groups’ data analysis (Fig.2C). All respiration features significantly varied between states One-way ANOVA Test (p<0.05) was first applied for global comparisons and then posthoc Tuckey (HSD) test for multiple comparisons (see legend Fig.2 for details).

Freq was particularly higher during AE (Fig.2C1) which was differentiated from QW by sniffing activity characterized by i) high iPF (Fig.2C2) and ePF (not shown) and ii) very short iDur (Fig.2C3) and eDur (not shown). iPF was much higher for waking states (AE and QW) than for sleep ones (SWS and REM). A slow Freq and a low iPF characterized sleep states. Similar observations and statistical differences were made on expiration features (not shown). To summarize, AE was characterized by a high iPF and a short iDur, whereas both sleep states exhibited a low iPF and a long iDur. QW regime appeared as unique with both iPF and iDur in the mid-range.

Then, the next question we asked was to what extent brain respiratory-drive, as described by others^3^, varied with brain state. We thus tracked respiration frequency in the LFP recordings of each structure, during each state. A simple visual inspection of power spectra (**Fig.3**) shows there is not a clear overlapping between respiration frequency and the low-frequency LFP oscillation, except during AE where the animals actively explored the environment. Nevertheless, it is difficult in this case to disentangle sniffing-related activity (6.2 ± 0.28 Hz) from the prominent exploration-related theta rhythm (about 7Hz, as measured in CA1 during REM, as proposed by Tort et al.). We observed that theta rhythm was a major frequency during all states, except during SWS where delta activity (0-5 Hz) increased. During QW, in addition to a clear theta rhythm in all structures (6.8 Hz) except OB (green line), spectral analysis evidenced a high power in the delta band (0-5 Hz), partly overlapping with respiratory frequencies. During SWS, while respiratory frequency was very slow and very constant around 1 Hz, LFP power was high in the whole delta band. Similarly, for REM state, respiration and LFP frequencies very poorly overlapped: while LFP power peaked in the theta range (around 7.5Hz), respiration frequencies lied in a much lower range (0.5-4Hz). To summarize, examination of the spectral characteristics of cerebral activity suggests that brain respiratory-drive probably varies with brain state. However, we did not observe a faithful reflection of the respiratory frequencies in the LFP of all areas so that it is difficult from power spectra to conclude on the existence of a respiratory drive of each brain structure.

**Figure 3:**
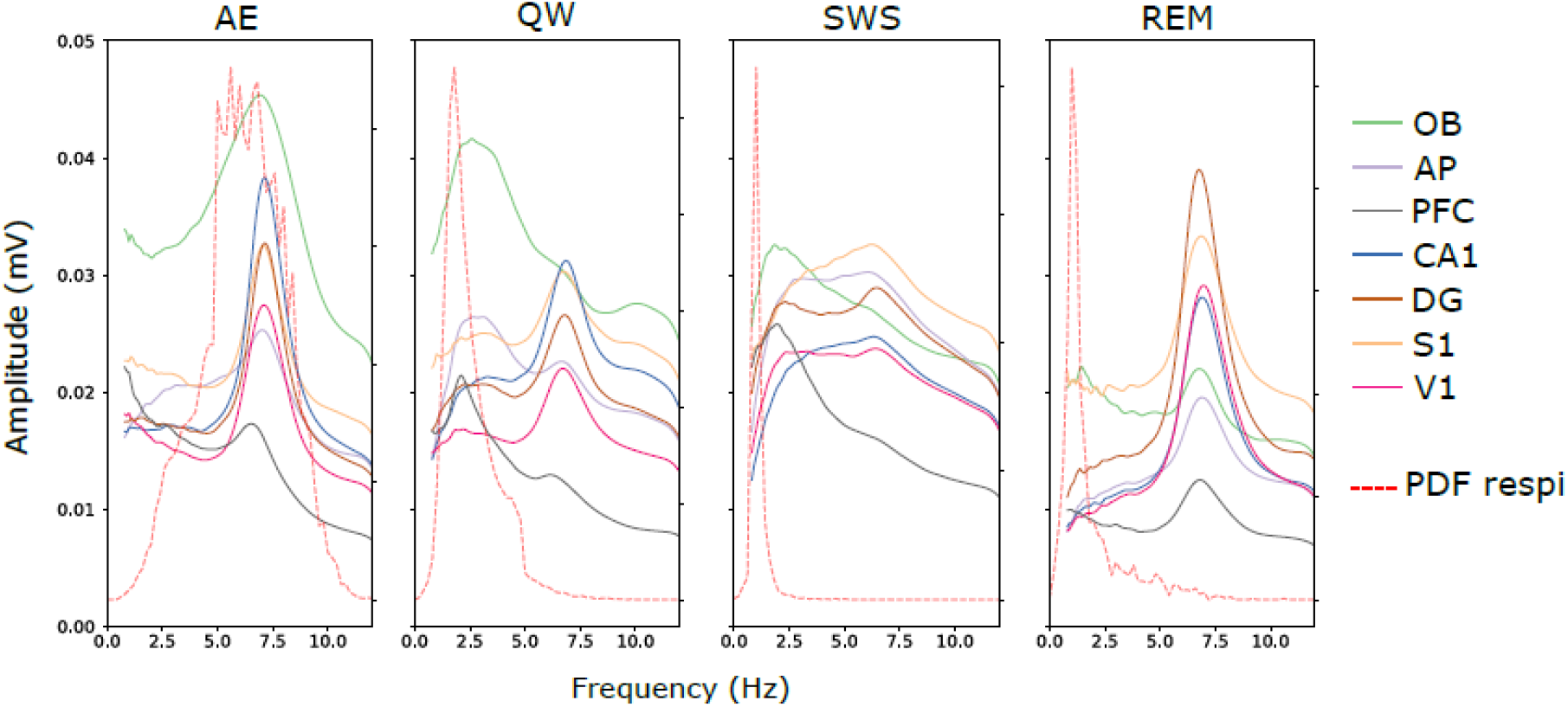
Power of LFP signals recorded in each structure (solid lines, in mV) along with the probability density function (PDF) of respiration frequencies (dotted red lines, no unit) during the four brain states (AE active exploration, QW quiet waking, SWS slow-wave sleep, REM rapid eye movements sleep). See sample size (number of rats) in Table 1.

To go further, we then analyzed the coherence between LFP signals and respiration in the low frequency band (0.5 – 10Hz). **Figure 4** shows LFP coherence to respiration for all the recorded brain regions during the four states. As described in Methods section, peak coherence values for actual data were compared to surrogates data. Actual and surrogates distributions significantly differed in all recorded regions and all states (see **Table Supp1**), suggesting that all the recorded areas were respiration-driven whatever the brain state. However, visual inspection of coherence spectra relativized this conclusion since it is obvious that coherence spectra are particularly flat during SWS and REM, whatever the structure. While extremely low during the sleep states, coherence appeared more clearly in waking states. During AE epochs, the respiration-LFP coherence was high (0.2 – 0.7) but only in olfactory structures (OB, AP) and PFC, and this on the whole range of Freq >5 Hz. It is noteworthy that QW state appeared as the only state during which coherence spectra appeared not flat in all the recorded areas. During SWS and REM, although power spectra exhibited a high power in the low frequency range, the coherence with respiration was very weak (10 times smaller than during QW), even if significant. To summarize, when analyzing respiration-LFP coherence over the whole respiratory frequency range, and not restricted to a specific frequency range, we observed that respiratory entrainment of brain areas varied with vigilance state, QW state being the state where LFP-respiration coherence is the highest in all the structures.

**Figure 4:**
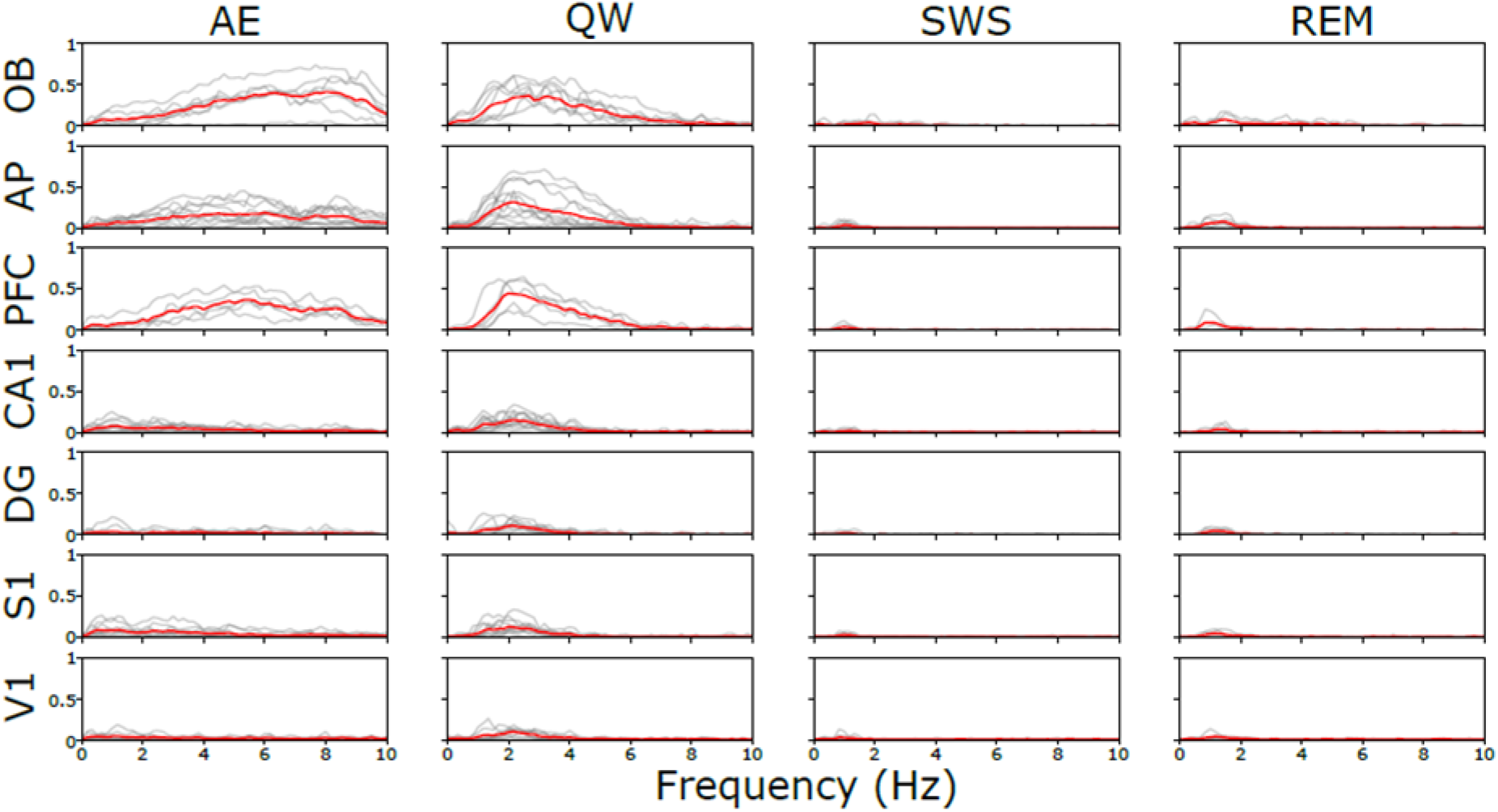
Coherence spectra between LFP and Respiratory signals during the four brain states (AE active exploration, QW quiet waking, SWS slow-wave sleep, REM rapid eye movements sleep), in all structures, in the 0.5-12 Hz range. Results obtained from simultaneous recordings in each rat (grey lines) and averaged across animals (red lines). See sample size (number of rats) in Table 1 and p values in Table Supp1.

Then, we searched in which conditions respiratory frequency could become the major frequency in the LFP. For this purpose, we computed covariation maps, as described in Methods, focusing on the major frequency expressed in LFPs in the 0.5-10 Hz range. This analysis beneficially complements coherence analysis since it gives access to cycle-by-cycle data. Results for all states and all structures are presented in **Fig.5**. A highlighted diagonal in the covariation maps indicates that LFP predominantly oscillated at the respiratory frequency. During AE, where rats sniffed in the 5-10Hz range, the dominant LFP frequency clearly followed respiration only in olfactory structures and interestingly in PFC (see the highlighted diagonal in OB, AP, PFC). Oppositely, covariation maps showed a uniform theta band activity whatever the respiration frequency (see the highlighted horizontal band in CA1, DG, S1, V1) indicating that these areas maintained an activity in the theta range whatever the respiration frequency. During QW, respiration frequency became the predominant frequency in LFP as assessed by a highlighted diagonal in the 1-4 Hz range. This clear covariation of respiration and LFP main frequency appeared in all the recorded structures. In four structures (S1, CA1, V1, DG) respiration frequency co-dominated with theta nevertheless (see the yellow spot around 7Hz). During SWS, the range of respiration frequencies was narrow (see the low variation in Fig.2C1) so that covariation analysis was not relevant. Whereas respiration frequency was slightly less constant (1.96±0.640 Hz) during REM, LFP predominant frequency was theta, except in the OB. We observed a covariation between respiration and LFP main frequency only in olfactory areas, PFC and DG. Finally, we found that LFP mainly oscillated at respiration frequency in all the recorded structures only during QW state.

**Figure 5:**
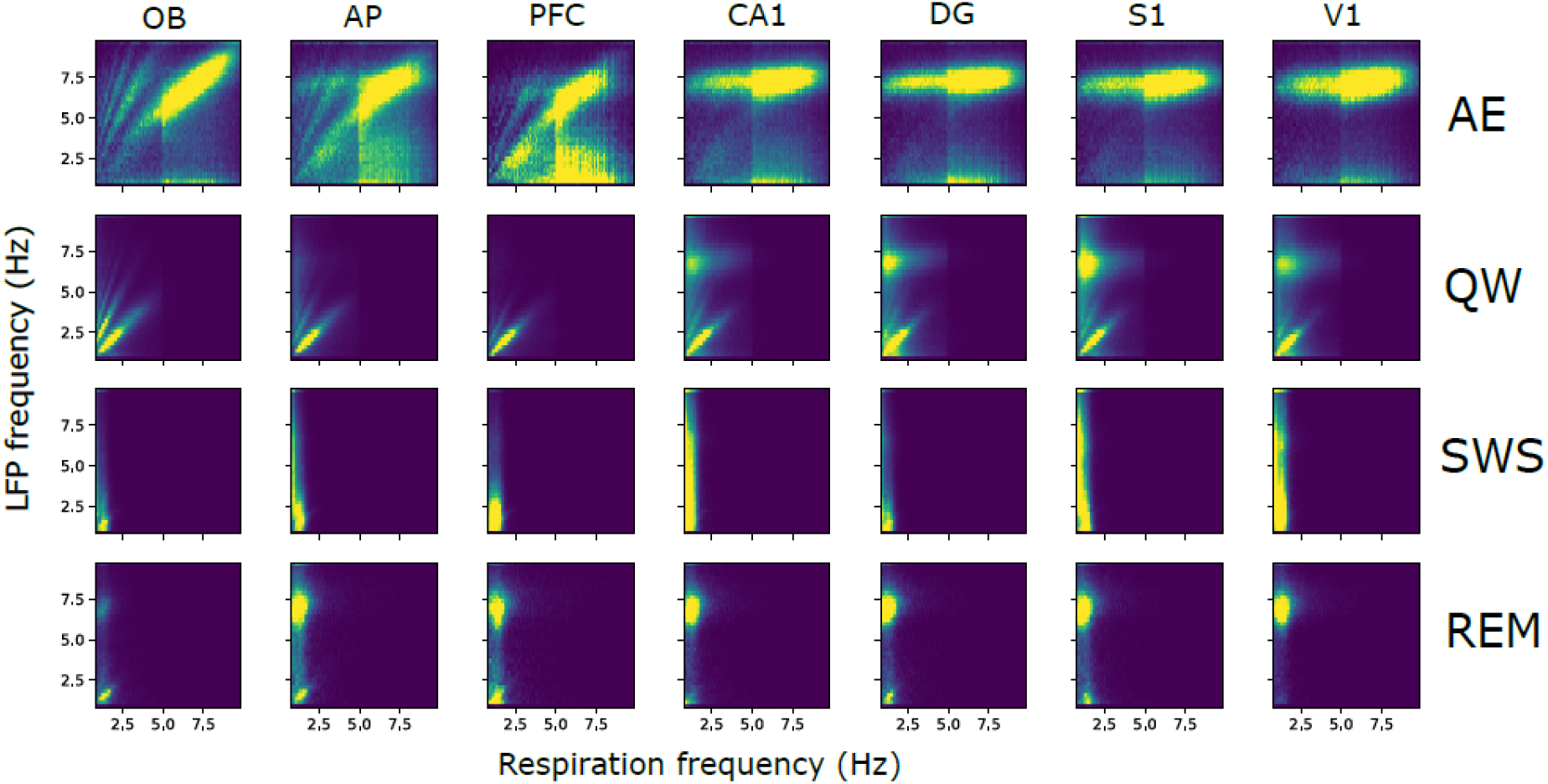
Covariation between LFP and respiration frequencies. Covariation maps obtained from LFP signals recorded under ambient air in OB, AP, PFC, CA1, DG, S1, and V1. Y-axis represents LFP frequency and X-axis respiratory frequency. The map is normalized so that the total sum is 1, and point density is represented on a color scale ranging from blue to yellow as the point density increases. See sample size (number of rats) in Table 1.

Respiratory drive of the brain is favored during the QW state but a question arises: is the specific respiratory entrainment of the brain during QW more related to the specificity of respiration features (frequency and/or peak flow rate) or to that of the QW state itself? To disentangle state *vs* respiration regime effect, we modified the respiration regime associated with each state by using a 5% CO_2_ enriched air in the plethysmograph in a subset of five rats. CO_2_ enrichment effectively strongly modified respiratory regime, as summarized in **Fig.Supp1.** Freq and iPF significantly increased with addition of CO_2_, except Freq during AE and REM. iDur significantly decreased except during QW, where it did not differed from ambient AIR condition. Covariation maps were computed for the same rats (n=5) submitted to both ambient AIR and CO_2_ conditions (**Fig.Supp2**). Under CO_2_ condition, covariation maps revealed changes (Fig.Supp2) that we objectivized by comparing the coupling index between respiration and the main LFP frequency in the different structures, during the four states (Fig.6). As explained in Methods, the coupling index is the sum of density along the diagonal of the covariation map and varies between 0 and 1 (1 is the highest coupling value). We did not perform any statistical test on these data because of the low sample size and we preferred showing individual data (Fig.6). During AE, the coupling index did not show much changes, except in olfactory areas and PFC. This probably indicates that, even if iPF increased significantly under CO_2_ (see Fig. Supp1), LFP frequency in these areas was always dominated by theta rhythm. During QW, the coupling index, which was already high under AIR condition, markedly increased, except in V1 where it increased very modestly. During SWS, even if delta band remained the major frequency, the coupling index increased under CO_2_ in most rats and most structures indicating that respiratory frequency became major more often than under AIR condition. We even observed highlighted diagonal on a short range of respiration frequencies (white arrows, Fig.Supp2B). During REM, while covariation maps exhibited a clearer and more extended highlighted diagonal (white arrows, Fig.Supp2B), the coupling index in Fig.6 appeared only exceptionally greater during CO_2_ than during AIR (1 rat in AP, all rats in PFC, 1 rat in DG, 2 rats in S1). This is probably because, even if respiratory frequency increased in the LFP signals, most often theta rhythm still dominated, as visible in the covariation maps of all structures except OB (Fig.Supp2B). Overall, these results suggest that the strong respiratory brain entrainment during QW was not related to the neuronal activity characteristics of this peculiar brain state but rather to the respiratory regime associated to this state. Indeed, when artificially extending the respiratory regime of the sleep states (with a larger band of frequencies and higher flowrates), respiration appeared as the major frequency in LFP signals where it was not observed with ambient air condition.

**Figure 6:**
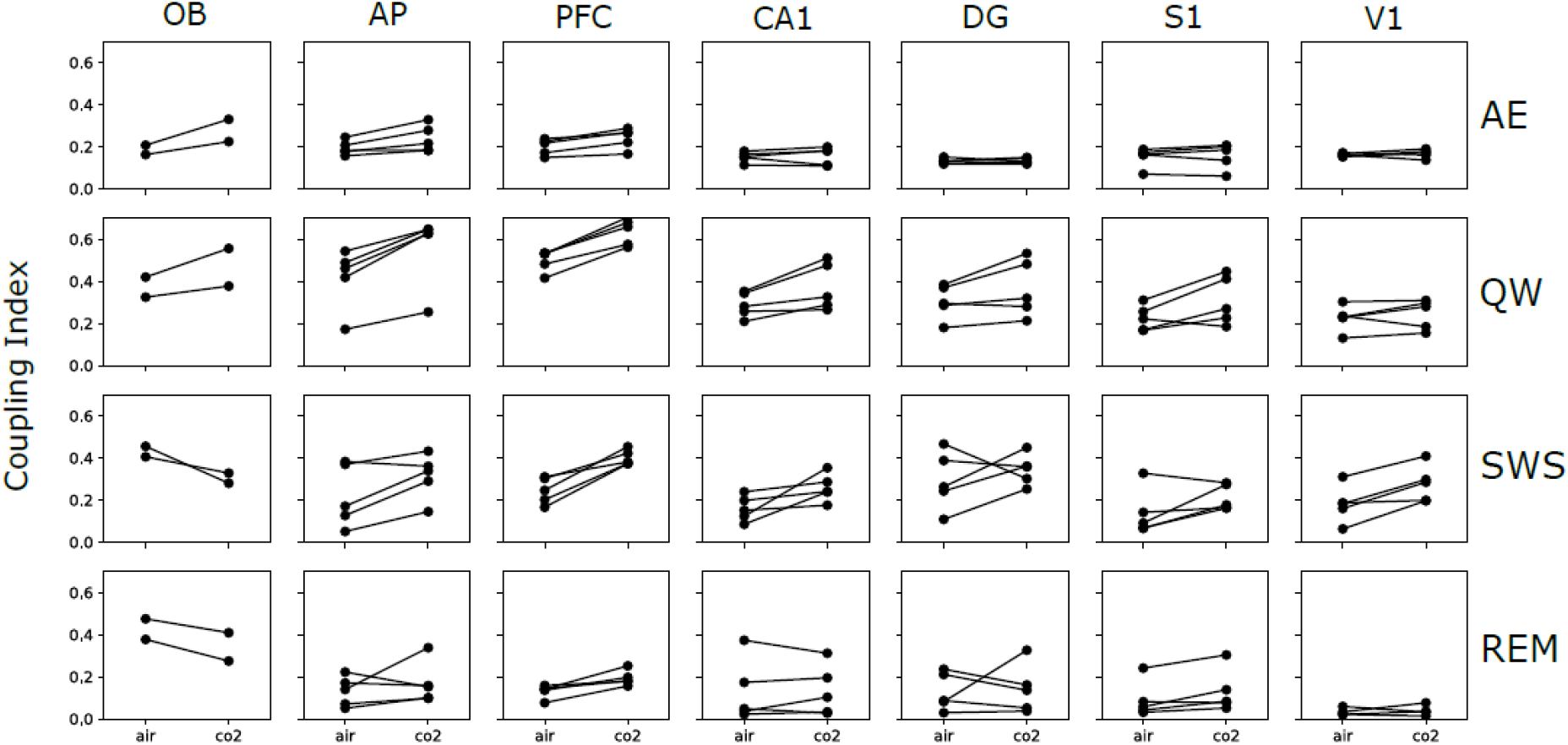
Comparison of respiration-LFP coupling between AIR and CO_2_ conditions. For each state (lines), each structure (columns), and each rat (black lines), the coupling index is plotted in AIR and CO_2_ conditions. AE active exploration, QW quiet waking, SWS slow-wave sleep, REM rapid eye movements sleep.

The next step was to decipher which of the respiration features was decisive in entraining brain structures to respiration since CO_2_ increased both Peak Flowrate and Frequency. To test this, we focused on two respiration features determining the deepness and frequency of respiration: inspiration peak flow rate (iPF) and inspiration duration (iDur). We then cycle-by-cycle analyzed the coupling between respiration and LFP frequencies, independently of the brain state, for the different values of iPF (**Fig.7A**) and iDur (**Fig.7B**). As visible both on the individual and averaged curves (sample size, see Table 1), the coupling index evolved following an inverted-U shape curve: almost null for low iPF values, it reached a maximum for values around 20mL.sec-1, then decreased for higher iPF values. This shape was similar in all structures, but with different amplitudes. Indeed, the respiration - LFP coupling as a function of iPF was very strong in olfactory structures and PFC, less so in other areas, but nevertheless always significant (quadratic term significance – i.e. existence of a local maximum - was assessed by comparing generalized mixed-effect linear models with Wald test (df = 1): OB: χ^2^=7.88, p=0.005; CA1: χ^2^=6.67, p=0.01; PFC: χ^2^=10.02, p=0.0015; DG: χ^2^=11.03, p=0.0009; AP: χ^2^=12.27, p=0.0004; S1: χ^2^=7.87, p=0.005; V1: χ^2^=7.88, p=0.005). Similar observations can be made when we analyzed the coupling index as a function of iDur (Fig.7B). As for iPF, the coupling index evolved following an inverted U-shape, optimal for a range of iDur values around 0.15sec (quadratic term was significant for PFC: χ^2^=6.34, p=0.01; and there were trends for AP: χ^2^=3.09, p=0.08; and OB: χ^2^=2.8, p=0.09). To explain the inverted-U shapes of the coupling index curves, we have to consider simultaneously the iPF and iDur. In Fig.7A, the highest values of iPF correspond to the sniffing cycles during AE, which present the shortest iDur. The fact that the curves go down beyond a certain range in Fig.7B (iDur>0.3sec) can be explained by the fact that the long iDur values correspond to the breathing cycles of sleep states, which also have small iPF values.

**Table 1:** Coordinates from bregma (mm) and number of rats (n) in each condition (Air and CO_2_)

| Brain structure | AP | ML | DV | n Air | n CO<sub>2</sub> |
| --- | --- | --- | --- | --- | --- |
| Olfactory bulb, OB | 5.5 * | 1.5 | 2 | 9 | 2 |
| Anterior piriform cortex, AP | 2.76 | 3.5 | ~6 | 13 | 5 |
| Dorsal hippocampus CA1, CA1 | -2.92 | 2 | ~3.5 | 13 | 5 |
| Dentate gyrus, DG | -4.8 | 2.8 | ~3.2 | 12 | 5 |
| Primary visual cortex, V1 | -5.28 | 4.2 | ~1.2 | 9 | 5 |
| Primary somatosensory barrel cortex, S1 | -2.4 | 5 | ~2 | 12 | 5 |
| Prefrontal cortex, PFC | 3 | 0.8 | ~3.5 | 5 | 5 |

\* from naso-frontal suture

**Figure 7:**
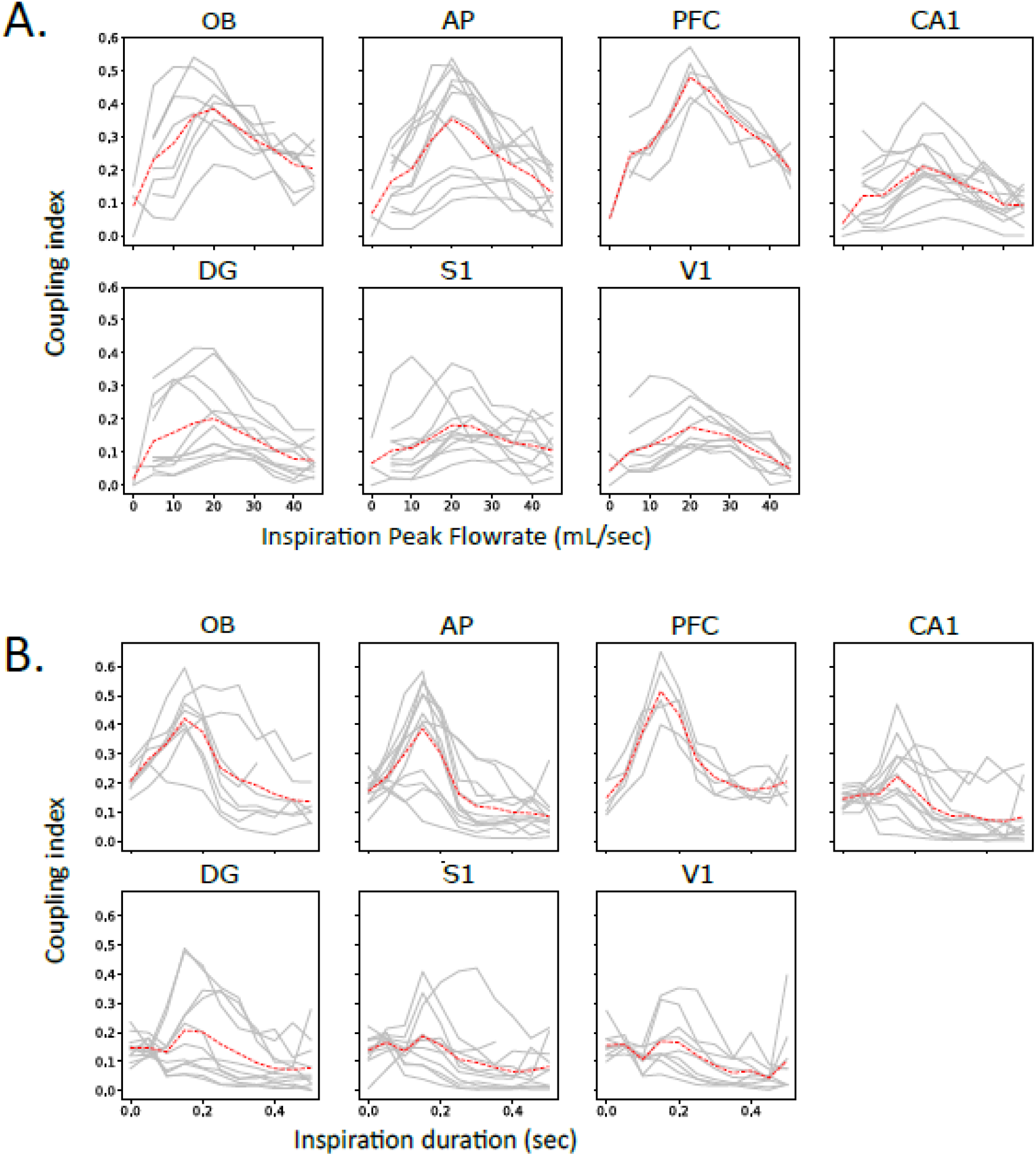
Coupling index as a function of inspiration peak flowrate iPF (**A**) and inspiration duration iDur(**B**). Data are presented individually for each animal (solid lines, see Table 1 for sample size), and averaged across animals (dotted red lines). The inverted U-shape of coupling index as a function of iPF and iDur was assessed by comparing two generalized mixed linear-models either with or without the quadratic term (see Methods for details). The comparison showed that the iPF quadratic term was significant for all areas (Wald test, p<0.02, see text for details) and that the iDur quadratic term was significant for some areas (Wald test, PFC: p=0.01, and trends for AP and OB, p<0.1, see text for details).

Finally, these results show that the best coupling between respiration and LFP frequencies is achieved when an inspiration peak flowrate in the range of 20-30mL/sec occurred simultaneously with a 0.1-0.2 sec inspiration duration. Such a trade-off between iPF and iDur corresponds to the specific features of respiratory regime associated with the QW state.

## DISCUSSION

We have systematically analyzed how respiration could drive brain areas during the spontaneous variations of respiratory regime related to four different behavioral states: active exploration, quiet waking, slow-wave sleep and paradoxical sleep. In addition to confirming the previous observations that LFPs in different brain structures can actually be respiration-driven^12–17^, we provided new fundamental data revealing that: 1) a large-scale respiratory drive is a specific characteristic of the quiet waking state, 2) this specificity is due to the respiratory regime associated with this state and not to the brain state itself, 3) an optimal respiratory regime for large brain entrainment is a deep and long inhalation (peak flowrate in the range of 20-30mL/sec and duration in the range of 0.1-0.2 sec).

We described for the first time that, even if each of the recorded structures can be occasionally entrained by respiration as reported by others^17,33^, a global respiratory drive of the brain only predominated during the quiet waking state. Before us, several groups described a respiratory drive in different brain areas with regard to the brain state. However, none of them reported this specificity of the quiet waking state. Indeed, Rojas-Libano et al. (2018) focused on periods of rest (our QW state) and exploration to investigate how gamma band activity in distant areas phase-locked to respiratory activity of the olfactory bulb. In an EEG study in the cat, Cavelli et al. (2019) observed the respiratory slow wave in the waking state but they did not differentiate rest from exploration. Tort et al. (2018) explored respiratory drive in different brain areas but, unlike us, chose to focus on two types of periods: exploration periods recorded in the animal home cage and REM sleep epochs recorded in a plethysmograph. Furthermore, because in both states theta oscillations and respiration may overlap in frequency, they also chose to analyze only epochs where respiration frequency and theta frequency were not overlapping. We took a different approach and explored LFP-respiration covariation i) during all the brain states, ii) during the whole range of respiratory frequencies. Finally, we have been the first to explore respiratory drive simultaneously in a large network of areas, during four brain states, and during the whole duration of recordings. This allowed us to highlight the specificity of the quiet waking state.

We also provided evidence that the specificity of quiet waking in allowing a global respiratory drive is not related to the state itself but rather to the respiratory regime associated. This demonstration has been possible because, using a CO_2_-enriched air in the plethysmograph, we modified the respiratory regimes associated with the different states. By extending the range of possible respiratory frequencies and amplifying nasal airflow, we have been able to observe that respiratory frequency could appear as the major LFP frequency during SWS and REM states, where it was not observable in ambient air conditions. We concluded that the global respiratory brain entrainment was determined by respiratory regime and not by the brain state itself. We chose to modify respiratory regime by CO_2_ rather than by stimulation of the respiratory nuclei^34^ principally because the method was simple to implement and was not invasive. However, CO_2_ use could be questionable because of its known effect on locus coeruleus^35,36^, on vegetative functions (increase in heart rate and respiratory rate), and on other complex behaviors such as anxiety^37^. Nevertheless, the very low concentration we used (5%) did not cause any significant change in the animals’ behavior. Once habituated to CO_2_ exposure, animals slept normally.

An interesting point in our study is our demonstration of an « optimal » respiratory regime for brain entrainment in rats, consisting in a deep (around 20mL.sec-1) and long (around 0.15 sec) inspiration. This demonstration has been made possible thanks to a cycle-by-cycle analysis that allowed us to study LFP-respiration coupling in details. We observed that a deep inspiration alone was not sufficient to entrain non-olfactory structures: if it was the case, we should have observed a large entrainment during active exploration where flowrates are the highest (see Fig.2). Indeed, during active exploration, inspiration was very ample but also very short. Similarly, a long inspiration alone was not sufficient to entrain non-olfactory areas since SWS and REM states exhibited the longest inspirations while entrainment was restricted to olfactory areas and PFC. During sleep sates, inspiration was very long but also very low in amplitude. Finally, the only state presenting an optimal breathing regime, with a deep and long inspiration, is the quiet waking state. We previously showed in the anesthetized tracheotomized rat that the respiratory drive of olfactory bulb depended on a trade-off between frequency and flowrate in the nasal cavity^30,32^. The present study extends this finding to non-olfactory areas and more importantly to the ecological condition of freely moving animal. Because brain respiratory-drive disappears when nasal respiration is by-passed or when activity of the olfactory bulb is suppressed^10,12,15,19,20^, it is tempting to conclude from our results that respiratory drive in the brain depends on the duration and flowrate with which respiratory airflows mechanically stimulate the olfactory epithelium^22^, thus activating the olfactory bulb. In such view, the olfactory bulb would represent the main driving force of the respiration influence to the brain. However, given that higher airflows are delivered by a deeper breathing and thus a different recruitment of respiratory centers and motor control, the effect of other pathways cannot be excluded.

Our study pointed out that all brain structures were not respiratory-driven with a similar probability. This has been also reported by Tort et al. (2018) showing that, in either REM or exploration, respiratory entrainment was most prominent in frontal regions (olfactory bulb, prelimbic cortex, anterior cingulate cortex). Among structures, PFC seems to be particularly sensitive to respiratory drive. Indeed, it is the only non-olfactory structure that appeared respiration-driven during the four brain states, whatever the respiratory regime. Individual data of LFP-respiration coupling index as a function of inhalation amplitude or duration (Fig.7) emphasized the strong and systematic coupling of this structure with respiration. PFC has been previously shown to be very well tuned to the 4 Hz respiratory frequency during animal freezing^38–41^. We show here, as we evoked in Dupin et al. (2019), that the strong tuning of PFC with respiratory rhythm holds whatever the respiratory frequency. This suggests that PFC could function as a hub structure, able to tune its activity to that of different areas according to the respiration frequency. Yet, PFC activity is coherent with basolateral amygdala during freezing when the animal breathes in the 2-4Hz range^38^, but it can also synchronize with hippocampal activity in the 7-8 Hz range^42–44^.

In their review, Heck et al. (2019) emphasized the potential role of respiration in organizing cortical activity in memory processes, notably by its ability to phase-lock hippocampal sharp wave ripples during wakening^15^. Our demonstration that brain structures are collectively and strongly susceptible to respiratory drive during quiet waking suggests that memory processes could be favored during this particular rest state. Our further demonstration that nasal airflow dynamics is crucial in the extent of brain entrainment by respiration could explain why memory consolidation is improved by nasal versus oral breathing^6^. The question now will be to test if an equivalent of the quiet state exists in human. We assume that such a state, characterized by a generalized entrainment of the brain on respiration, could be reached during deep-breathing practice.

The crucial role of the olfactory pathway in brain respiratory-drive is today largely recognized, and particularly well emphasized in review papers^2,45–48^. But credit where credit is due: in their excellent opinion paper, Fontanini and Bower^49^ were the first in 2006 to clearly posit that *“[…] it might be that the pan-cortical correlations that particularly characterize slow-wave sleep and meditation actually depend on an afferent input signal generated within the olfactory system*.” Our work is definitely in line with this thinking and that of a “sniffing brain” they developed. Moreover, our new data make it possible to answer in the affirmative to the question Heck et al. (2019) asked in their recent review: “*Does the change in sniff frequency (for rats, rabbits, cats, and mice) or duration (for humans) alter the cortical circuit by driving the OB differently via the olfactory nerve […]*”. Finally, our results suggest that nasal airflows adjustment by variation in respiratory regimes could be an effective tool for influencing brain activity. Notably, the slow and deep breathing associated to rest state seems to be an optimal regime to achieve a respiratory drive on the whole brain.

## METHODS

### Experimental Procedures

#### Animal care

The 13 male Long Evans rats (Janvier Laboratories, Le Genest Saint Isle, France; 8 weeks old, ~250-300 g at the start of the experiment) used were housed in groups of four in a temperature (22±1 °C) and humidity (55±10%) controlled room and exposed to a 12/12 h light/dark cycle (light onset, 6:00 am). Experiments were conducted during the light period (between 9:00 am and 3:00 pm). Food and water were available *ad libitum*. All experiments were performed in accordance with Directive 2010/63/EU of the European Parliament and of the Council of the European Union regarding the protection of animals used for scientific purposes (National Ethics Committee, Agreement APAFIS #17088).

#### Animal preparation

Preparation consisted in electrode implantations in different brain areas. Anesthesia was induced by an initial dose of a mixture of chloral hydrate and sodium pentobarbital (intraperitoneal; Equithesin 3 ml/kg) and maintained with additional doses as needed. Rats were administered with anti-inflammatory and analgesic treatment (subcutaneous; carprofen 2 mg/kg or meloxicam 0.2 mg/kg) immediately after surgery and during several postoperative days if necessary. After a minimal 10 days’ period of recovery, animals were implanted with monopolar stainless steel LFP recording electrodes (diameter: 100 μm, 200-800 kΩ impedance, California Fine Wire) soldered to a copper wire (diameter: 250 μm). Electrodes were positioned stereotactically into the left cortical hemisphere in 6 (8 rats) or 7 (5 rats) brain areas (see Table 1 for coordinates and sample size) comprising the olfactory bulb (OB), anterior piriform cortex (AP), dorsal part of the hippocampus CA1 (CA1) and dentate gyrus (DG), primary visual cortex (V1), primary somatosensory barrel cortex (S1), and prefrontal cortex (PFC). Electrophysiological activity was recorded while the electrode was lowered. Positioning of the recording electrode in the AP was determined by the shape of evoked field potential induced by stimulation of a bipolar OB electrode placed close to the mitral cell layer. The final location was selected to where the evoked potentials had the largest amplitude (before reversal). The stimulation electrode was then replaced by a monopolar recording electrode in the OB. The depth of the recording electrodes in the OB and CA1 was adjusted using their large multiunit activity, to the level of the mitral cell layer and just below the pyramidal layer, respectively. Positioning of the other recording electrodes tips (DG, S1, V1, PFC) was achieved stereotactically. Each electrode was individually fixed to the skull using dental cement. A reference wire was connected to a skull golden screw located above the posterior portion of the contralateral cortical hemisphere. Two anchor screws were also inserted in the contralateral side to secure the implant. Each electrode was attached to a 32-pin electrode interface board (EIB, NeuraLynx, Inc, USA, ViewPoint France) combined with an omnetics connector and centered on the animal’s head.

#### LFP and Respiration recording

Signals were acquired by telemetry using a 32 channels wireless recording system (W32 headstage, TBSI, ViewPoint France). Signals were sampled at 15 kHz, amplified (gain 800x) and recorded via an acquisition card (USB-2533, Measurement Computing, Norton, MA). Signals were acquired using custom-made open-source software (pyacq, https://github.com/pyacq/pyacq) and stored on a computer for offline analysis.

Respiratory activity was recorded by using whole-body plethysmography. Plethysmograph (EMKA Technologies, France) was previously described in detail^50,51^.

One camera (B/W CMOS PINHOLE camera) was placed in a corner of the cage in order to monitor the animal behavior.

#### Recording sessions

Before starting recording sessions, animals were habituated to the plethysmography cage during 2 sessions of 20 min. At the end of this period, animals were completely familiarized with the environment and did not show any sign of stress.

##### Ambient air

Recording sessions then started. Six sessions (one session per day) of 2 hours under ambient air were recorded for each animal. During the first minutes, animals alternated between active exploration and quiet rest. After an amount of time, varying between animals (10-30 min), rats started to sleep, with SWS and REM states alternation.

##### CO_2_-enriched air

In some experiments, we used a 5% CO_2_-enriched air in the plethysmograph in order to change respiration regime. The choice of 5% concentration was based on respiratory physiology experiments, where the CO_2_ concentration usually used is 8%^52^. It has been shown that a concentration of 10% can increase immobility bouts in Long-Evans Rats, suggesting a certain level of anxiety^53^. We therefore searched for the lowest concentration able to induce a change in the amplitude and/or respiratory frequency. Our criteria were that: 1) animal could fall asleep with the same delay as under ambient air, 2) no signs of panic (no increase in episodes of immobility or grooming) were shown, 3) sniffing frequency returned to baseline in less than 5 minutes after CO_2_ stopping in the plethysmograph. Pilot experiments on 4 test animals (not shown) allowed us to define 5% CO_2_ concentration as obeying all of these criteria. Following the 3 ambient air recording sessions, five animals were recorded during 3 new sessions under CO_2_-enriched air (one session per day). To diffuse CO_2_ enriched air, a 10% CO_2_ – air container (Air Liquide, France) was connected to the plethysmograph. Pure air was mixed with CO_2_ (50/50) so that the final concentration was 5%. Under CO_2_, the recording sessions were limited to 2 hours. Exposure of animals 3 times 2 hours was not sufficient to induce chronic hypercapnia^54^.

### Data analysis

#### Respiratory signal

Respiratory signals were acquired and extracted using previously described method^51^ allowing accurate measurements of different respiratory parameters in behaving animals. The detection of the respiratory cycles was achieved using an enhanced version of the algorithm described in a previous study^55^. This algorithm performs two main operations: signal smoothing for noise reduction, and detection of zero-crossing points to define accurately the inspiration and expiration phase starting points. For each respiratory cycle, inspiration and expiration peak flowrate (iPF, ePF), inspiration duration and expiration duration (iDur, eDur) were measured. Instantaneous respiratory frequency (Freq) was determined as the inverse of the respiratory cycle (inspiration plus expiration) duration.

#### States scoring

Based on video, electrophysiological, and breathing recordings, two well-trained experimenters visually coded four vigilance states by inspection of behaviors, respiration, and associated LFP spectral features (see Fig. 1). State was classified as active exploration (AE) if the animal engaged in exploratory behavior (locomotion, whisking, and sniffing), with a high amplitude and >5Hz respiration^33^ during at least 3 cycles, and a cortical LFPs expressing high gamma (30–55 Hz) power density. In quiet waking (QW), the animal was immobile (standing or sitting quietly) with a high amplitude and <5Hz respiration, cortical LFPs expressed relatively high gamma activity. In slow wave sleep (SWS), the animal was lying immobile with eyes closed and slow and extremely regular respiratory movements. LFP were characterized by prominent delta waves (1–4Hz). In rapid eye movements sleep (REM), the animal was immobile with eyes closed. Breathing was irregular, with epochs of low and slow amplitude alternating with short epochs of higher frequency and amplitude (see Figs. 1 & 2). LFP were low in amplitude and expressed very high theta and gamma power. Intermediary states, where LFP features were not very clear, were not coded.

#### Electrophysiological signals

Data were analyzed using custom-written scripts with the python language and its scientific ecosystem tool suite.

##### Spectral analysis

The power spectral density (PSD) of the LFP signals was calculated using the continuous Morlet wavelet transform^56^ instead of the classical windowed Fourier transform (FFT). The Morlet wavelet estimated the amplitude of the signal across time and frequency. The obtained time-frequency map was then segmented in periods of interest with variable durations and averaged on the time axis. We made this choice mainly because low frequency analyses require very long windows with FFT methods while the Morlet wavelet transform is a continuous representation along time. Segmenting state periods with varying durations combined with fixed windowed method would be sub-optimal whereas the continuous representation in the Morlet time frequency map makes the process easier and more accurate. The drawback with our approach is that the final PSD is smoother on the frequency domain than a standard FFT.

##### Coherence analysis

For the computation, the signal was down sampled to 1000Hz. LFP-respiration coherence was calculated with a custom-modified version of Scipy coherence function. Indeed, since coherence is based on FFT window method then the outcome is biased by the total duration of the period. In our case, to compensate the great difference in duration between states, we randomly sub-sampled an equal number of non-juxtaposed epochs of each state and then used the classic coherence with no overlap on these epochs (256 epochs of 6s). Coherence between LFP and respiration was compared against chance using a surrogate-based statistical testing. Surrogate were obtained by preserving the LFP and respiration spectra using a 4-steps process: 1) shuffling (n=500) of 256 epochs, 2) coherence computing, 3) extraction of the maximum coherence value, and 4) computation of the distribution of surrogate values. Finally, the actual values of the maximum of coherence spectrum was compared to this surrogate distribution providing an estimation of p-value.

##### Covariation maps and Coupling Index

To study frequency-frequency coupling between LFP signals and respiration, we designed a homemade method that allowed tracking instantaneous frequency synchrony^41^. To do so, for each detected respiratory cycle, the frequency was estimated as 1/cycle duration and the time course of the instantaneous respiratory frequency was extracted. In parallel, the continuous Morlet scalogram for the LFP signal was computed in our frequency band of interest (0.8–10 Hz). In each time bin (4 ms), we extracted the frequency of the point of maximum instantaneous power. Finally, this sequence of frequencies gave us the time course of the LFP predominant instantaneous frequency. From the two times series obtained (instantaneous respiration frequency and predominant instantaneous LFP frequency), a 2D matrix histogram was built, with the respiratory frequency represented on the x-axis and the LFP frequency on y-axis. This 2D histogram, called covariation map, was normalized so that the total sum equaled 1, and point density was represented on a color scale ranging from blue to yellow as the density of points increased. A coupling between respiration frequency and LFP frequency can be assumed when a high density (i.e., yellow color) was observed along the diagonal of the covariation map (see Fig. 5 for an illustration). Conversely, a non-correlated Gaussian shape indicated that the frequency of respiration and the main frequency of LFP varied independently. The two possibilities can co-occur on the same covariation map.

Based on this map, we were then able to introduce a new metric: the sum of density along the diagonal. To compute the sum of the diagonal, we first applied a smoothing Gaussian kernel (*σ*= 0.58 Hz) to assess the density around the diagonal. This metric estimated the coupling strength between respiration and LFP with a factor in 0-1 range where 1 indicates that respiration frequency is always the dominant LFP frequency and 0 signals the two frequencies are never equal. We named “coupling index” this custom metric. To validate the accuracy of this metric, we constructed surrogates (n=500) by applying random time-shift between the respiration time series and LFP time series. This procedure allows to remove the instantaneous coupling but maintains the global frequency distribution of both time series. The measured coupling index was always higher than the 0.9999 quantile of the surrogate distribution.

The two time-series (respiration frequency and LFP frequency) describing the covariation map can be segmented in short periods of variable duration before computing the covariation map. Therefore, the frequency time series are segmented on a respiratory cycle-by-cycle basis. This segmentation allows to group cycles with common characteristic, for instance iPF or iDur. For each of these groups of cycles, we can compute a covariation map and the corresponding coupling index. Ultimately, we were able to display a coupling index vs “inspiration peak flow rate” (or duration) plot across animal state and on an animal-by-animal basis. In some cases, a total duration of 60sec was not reached: the missing values were then not considered for the computation.

While looking at latter analyses, we often noticed the existence of a local maximum of coupling index for intermediate values of a given respiratory cycle characteristic. In order to statistically assess the existence of this intermediate maximum, we fitted our data with generalized linear mixed-effect models. Model fixed-effects included a first or second order polynomial of the respiratory cycle characteristic, while random effects always included a second order polynomial. Data were grouped per animal (distinct models have been done for each area). The link function was a logarithm and the variance described with a Gamma function (family function). We visually checked homoscedasticity and normality of the residuals after model fitting. Comparison between nested models was obtained with a Wald chi-square test. Mixed-model analysis was done with software R 3.4.4 and library lme4 1.1-21.

## Data availability

All data and scripts at any stage of analysis are available upon reasonable requests from Nathalie Buonviso.

## Acknowledgements

This work was supported by the Centre National de la Recherche Scientifique and the LABEX CORTEX (ANR-11-LABX-0042) of Université de Lyon, within the program “Investissements d’Avenir” (ANR-11-IDEX-0007) operated by the French National Research Agency. We thank Emmanuelle Courtiol and Anne-Marie Mouly for careful reading of this manuscript.

