## Supplementary material for "The deep and slow breathing characterizing rest favors brain respiratory-drive"

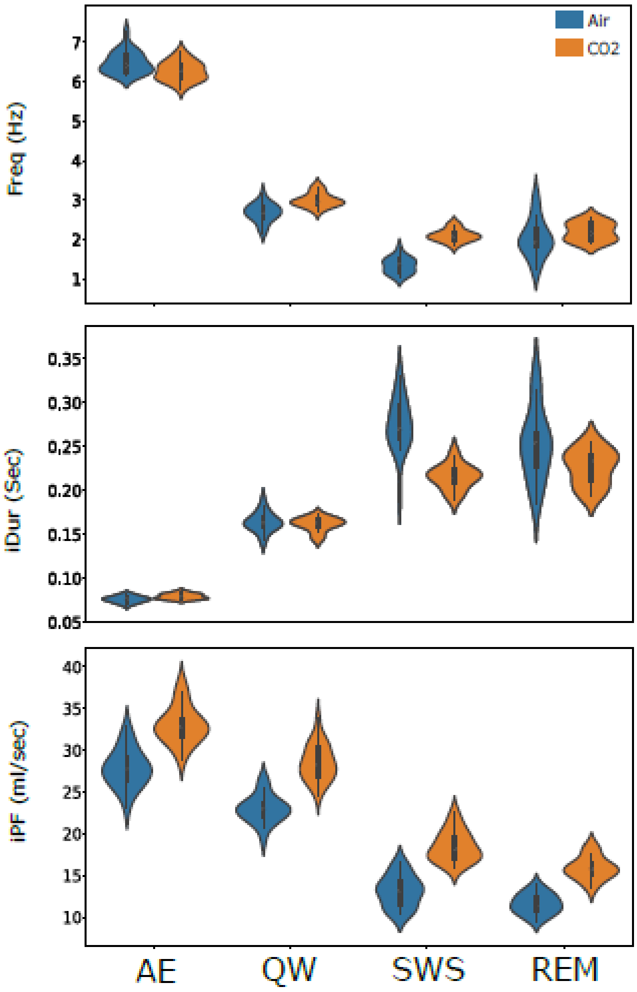


| p-value | Freq | iPF | iDur |
|---|---|---|---|
| AE | 0,008 | <0,001 | 0,001 |
| QW | <0,001 | <0,001 | 0,357 |
| SWS | <0,001 | <0,001 | <0,001 |
| REM | 0,234 | <0,001 | 0,026 |

**Fig.Supp1**: Difference of respiration frequency (Freq, **top panel**), inspiration duration (iDur, **middle panel**) and inspiration peak flowrate (iPF, **bottom panel**) between ambient AIR and CO2-enriched conditions. **Table**: Each parameters of each state (AE active exploration, QW quiet waking, SWS slow-wave sleep, REM rapid eye movements sleep) are compared between ambient AIR and $CO_2$- enriched air using a T-test.

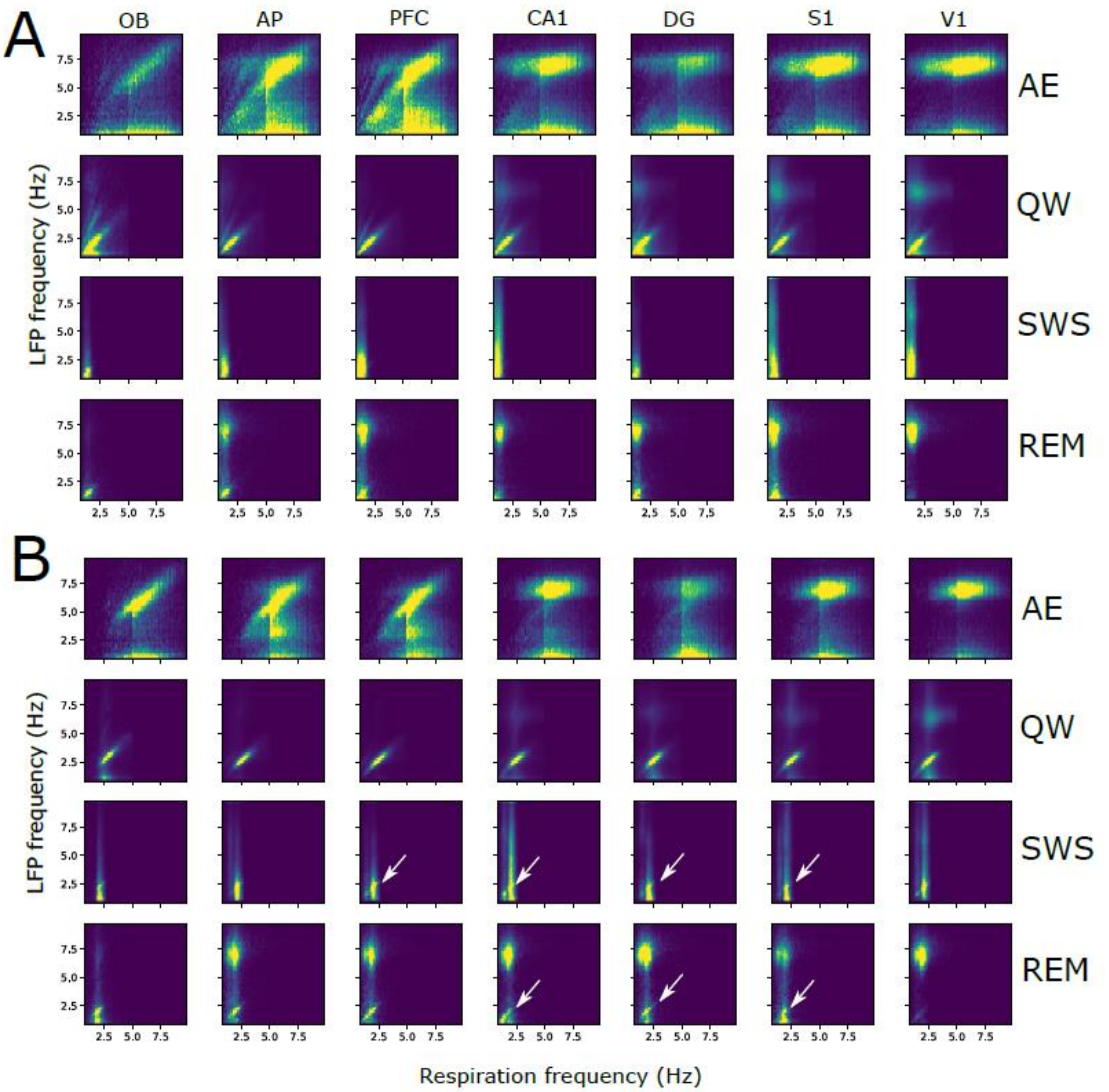


**<u>Fig.Supp2</u>**: Covariation between LFP and respiration frequencies. Covariation maps obtained from LFP signals recorded under ambient air (**A**) and $CO_2$ (**B**) conditions in OB, AP, PFC, CA1, DG, S1, and V1. Y-axis represents LFP frequency and X-axis respiratory frequency. The map is normalized so that the total sum is 1, and point density is represented on a color scale ranging from blue to yellow as the point density increases. See Table 1 for samples size. AE active exploration, QW quiet waking, SWS slow-wave sleep, REM rapid eye movements sleep.

**Table Supp1**: LFP-respiration coherence. For each state (AE, QW, SWS, REM) and each structure (OB, AP, PFC, S1, CA1, V1, DG), the peak of coherence values (Frequency) between LFP and respiration were compared between actual and surrogates data using the process described in Methods. Are indicated in red and orange those conditions were coherence in actual data were ≥ 0.3 or 0.1 respectively.

| State | Structure | Frequency | actual | surrogate |
|---|---|---|---|---|
| AE | OB | 8.00 | 0.41 | 0.000 |
| | AP | 6.00 | 0.19 | 0.000 |
| | PFC | 5.33 | 0.37 | 0.000 |
| | CA1 | 1.17 | 0.08 | 0.000 |
| | DG | 1.00 | 0.04 | 0.001 |
| | S1 | 1.17 | 0.09 | 0.000 |
| | V1 | 1.17 | 0.05 | 0.000 |
| QW | OB | 2.67 | 0.37 | 0.000 |
| | AP | 2.17 | 0.32 | 0.000 |
| | PFC | 2.00 | 0.45 | 0.000 |
| | CA1 | 2.17 | 0.15 | 0.000 |
| | DG | 2.17 | 0.11 | 0.000 |
| | S1 | 2.00 | 0.13 | 0.000 |
| | V1 | 2.17 | 0.10 | 0.000 |
| SWS | OB | 1.83 | 0.05 | 0.000 |
| | AP | 1.00 | 0.04 | 0.000 |
| | PFC | 1.00 | 0.04 | 0.000 |
| | CA1 | 1.17 | 0.02 | 0.005 |
| | DG | 1.17 | 0.02 | 0.007 |
| | S1 | 1.00 | 0.03 | 0.001 |
| | V1 | 0.83 | 0.03 | 0.001 |
| REM | OB | 1.50 | 0.08 | 0.000 |
| | AP | 1.50 | 0.08 | 0.000 |
| | PFC | 1.00 | 0.10 | 0.000 |
| | CA1 | 1.33 | 0.05 | 0.000 |
| | DG | 1.17 | 0.05 | 0.000 |
| | S1 | 1.17 | 0.05 | 0.000 |
| | V1 | 1.17 | 0.04 | 0.000 |
